# The pathobiology of *Mycobacterium abscessus* revealed through phenogenomic analysis

**DOI:** 10.1101/2021.10.18.464689

**Authors:** Lucas Boeck, Sophie Burbaud, Marcin Skwark, Will H. Pearson, Jasper Sangen, Aaron Weimann, Isobel Everall, Josephine M Bryant, Sony Malhotra, Bridget P. Bannerman, Katrin Kierdorf, Tom L. Blundell, Marc S. Dionne, Julian Parkhill, R. Andres Floto

## Abstract

The medical and scientific response to emerging pathogens is often severely hampered by ignorance of the genetic determinants of virulence, drug resistance, and clinical outcomes that could be used to identify therapeutic drug targets and forecast patient trajectories ^1–5^. Taking the newly emergent multidrug-resistant bacteria *Mycobacterium abscessus* as an example ^6^, we show that combining high dimensional phenotyping with whole genome sequencing in a phenogenomic analysis can rapidly reveal actionable systems-level insights into bacterial pathobiology. Using *in vitro* and *in vivo* phenotyping, we discovered three distinct clusters of isolates, each associated with a different clinical outcome. We combined genome-wide association studies (GWAS) with proteome-wide computational structural modelling ^7^ to define likely causal variants, and employed direct coupling analysis (DCA) ^8^ to identify co-evolving, and therefore potentially epistatic, gene networks. We then used *in vivo* CRISPR-based silencing to validate our findings, defining a novel secretion system controlling virulence in *M. abscessus*, and illustrating how phenogenomics can reveal critical pathways within emerging pathogenic bacteria.

## INTRODUCTION

Over the last two decades, *M. abscessus*, a rapid growing species of nontuberculous mycobacteria (NTM), has emerged as a major threat to individuals with Cystic Fibrosis (CF) and other chronic lung disease ^9^. Rates of infection of CF patients have increased around the World ^9–11^, in part due to hospital-based person-to-person transmission ^12,13^ and the emergence of globally-spread dominant circulating clones that are associated with increased virulence and worse clinical outcomes ^14^. Infections with *M. abscessus* are challenging and sometimes impossible to treat ^9,15,16^, lead to accelerated inflammatory lung damage ^17,18^, and may prevent safe transplantation ^19^.

To date, very little is known about how *M. abscessus* infects humans ^6^, how it causes inflammatory lung damage, and how it resists antibiotics ^6^. There is thus an urgent need to better understand the pathophysiology of *M. abscessus*, to define optimal drug targets, and to predict the virulence and antibiotic susceptibility of clinical isolates.

We therefore sought to combine detailed *in vitro* and *in vivo* phenotyping, whole genome sequencing, computational structural modelling, and epistatic analysis to provide a phenogenomic map of *M. abscessus* that might define critical pathways involved in virulence and drug resistance.

## RESULTS

We first characterised 331 clinical *M. abscessus* isolates across 58 phenotypic dimensions exploring five key pathogenic traits: planktonic growth in different carbon sources; antibiotic resistance (at early and late time points) against a selection of drugs recommended by clinical treatment guidelines ^9^; *in vitro* infection of a human macrophage cell line model (differentiated THP1 cells), monitored using high content confocal microscopy; *in vivo* infection of *Drosophila melanogaster*, measuring host survival and inflammatory responses; and clinical outcomes following infection, available through previously collected metadata ^14^ (***Figure 1a, Supplementary Figure 1***).

**Figure 1:**
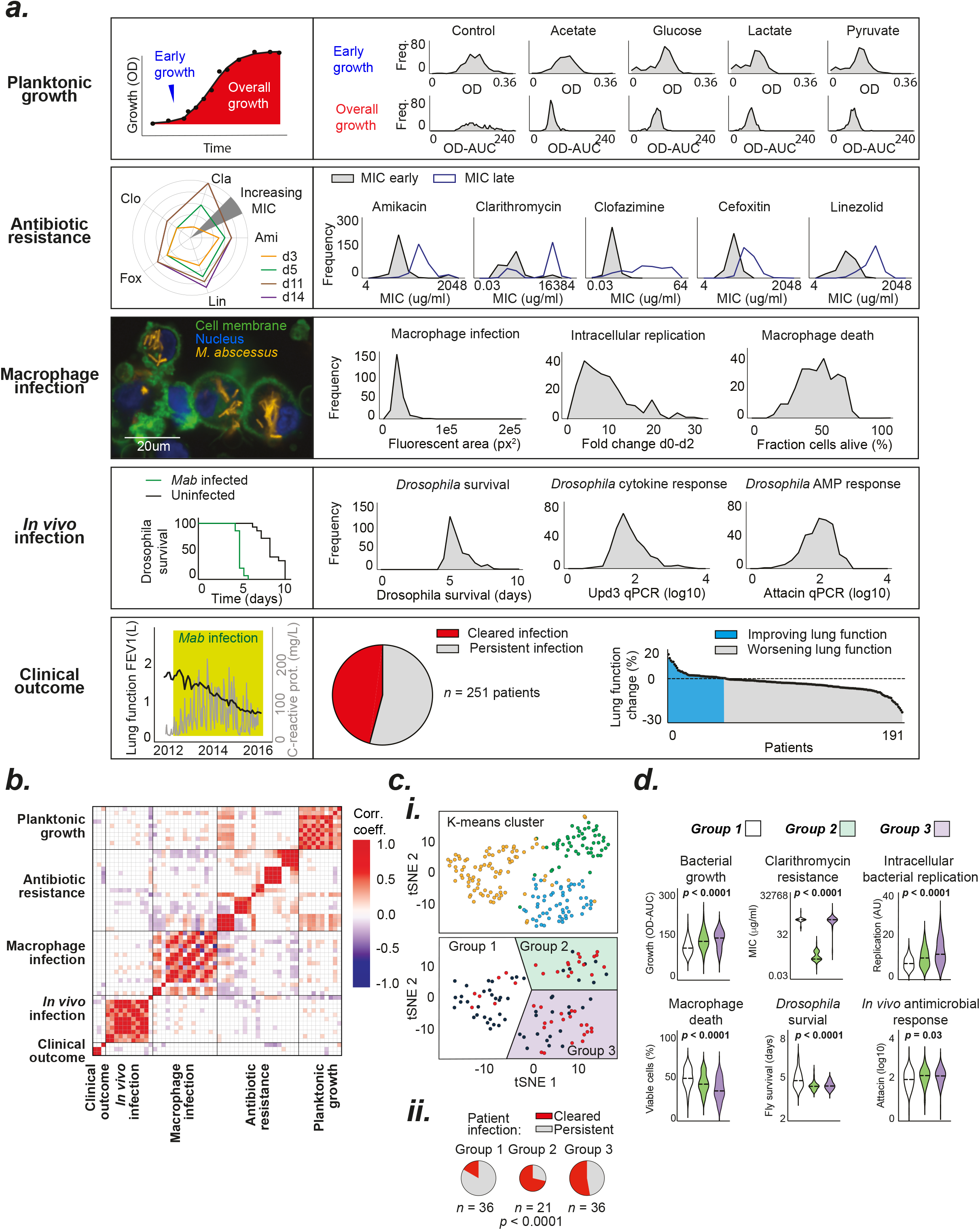
Multidimensional phenotyping of *M. abscessus*. (**a**) Phenotypic variability of clinical *M. abscessus* isolates was assessed across multiple dimensions (described in Supplementary Methods) including: (i) Planktonic growth (assessed by serial OD measurement) in a range of different carbon sources; (ii) Minimal inhibitory concentrations (MIC) of a range of clinically relevant antibiotics were assessed on day 3 (MIC early) and day 11 (MIC late) to quantify intrinsic and inducible drug resistance; (iii) Macrophage infection (4h post infection), intracellular replication (2d post infection) and death (2d post infection) were quantified using high-content imaging of differentiated THP-1 cells incubated with tdTomato-expressing clinical isolates; (iv) Survival and immune response of *Drosophila melanogaster* infected with clinical isolates; and (v) Clinical outcomes (lung function decline and clearance of *M. abscessus* from sputum samples) of infected patients (**b**) Pearson correlations within and across phenotypic groups shown as a matrix, with non-significant (p > 0.05) associations shown in white. (**c**) Clustering of clinical isolates, using k-means and T-stochastic neighbour embedding, based on (i) experimentally observed phenotypes only, demonstrate three distinct groups that (ii) differ in their clinical outcomes. (**d**) Distribution of specific phenotypes across the three phenotypic groups. P values calculated using Chi-squared test or one-way analysis of variance, as appropriate.

We examined the relationship between phenotypes, finding correlations within, and sometimes between, pathogenic traits (***Figure 1b, Supplementary Figure 2***). To explore whether there were distinct patterns of bacterial behaviours, we used experimentally-derived data to plot individual isolates in phenotypic space, identifying three discrete groups, each associated with different clinical outcomes (***Figure 1c,d, Supplementary Figure 3***), representing distinct heritable traits (***Supplementary Figure 4***). Isolates in Groups 1, 2, and 3 demonstrated progressively faster growth in liquid culture and within macrophages, higher rates of macrophage and *Drosophila* death, and greater inflammatory responses. While Group 2 isolates were associated with the most favourable clinical outcome, potentially related to their macrolide susceptibility (a key determinant of treatment response ^20,21^), we found that Group 1 and Group 3 isolates, although similarly macrolide resistant, had very different clinical outcomes, highlighting the importance of other phenotypic characteristics in determining prognosis, and suggesting that more virulent (and thus immunogenic) isolates might be cleared more easily by patients (as previously reported for other pathogenic bacteria ^22–25^).

To understand the genetic basis for these important variations in *M. abscessus* behaviour, we used whole genome sequence data to perform a genome-wide association study (GWAS) for each phenotype (***Figure 2a***), evaluating approximately 270,000 genetic variants comprising single nucleotide polymorphisms (SNPs), insertions, and deletions. We used mixed models corrected for population structure ^26^ to identify locus effects, as well as uncorrected linear models to ensure we captured lineage effects ^27^. In total, we identified 1926 hits (involving 1000 genes) across 46 phenotypes (***Supplementary Table 1***). These included previously known genetic determinants, such as the 16S and 23S rRNA mutations associated with constitutive aminoglycoside and macrolide resistance (*p* = 1.3 x 10^-75^ and *p* = 1.5 x 10^-54^ respectively; ***Supplementary Figure 5***), thereby confirming the effectiveness of our approach.

**Figure 2:**
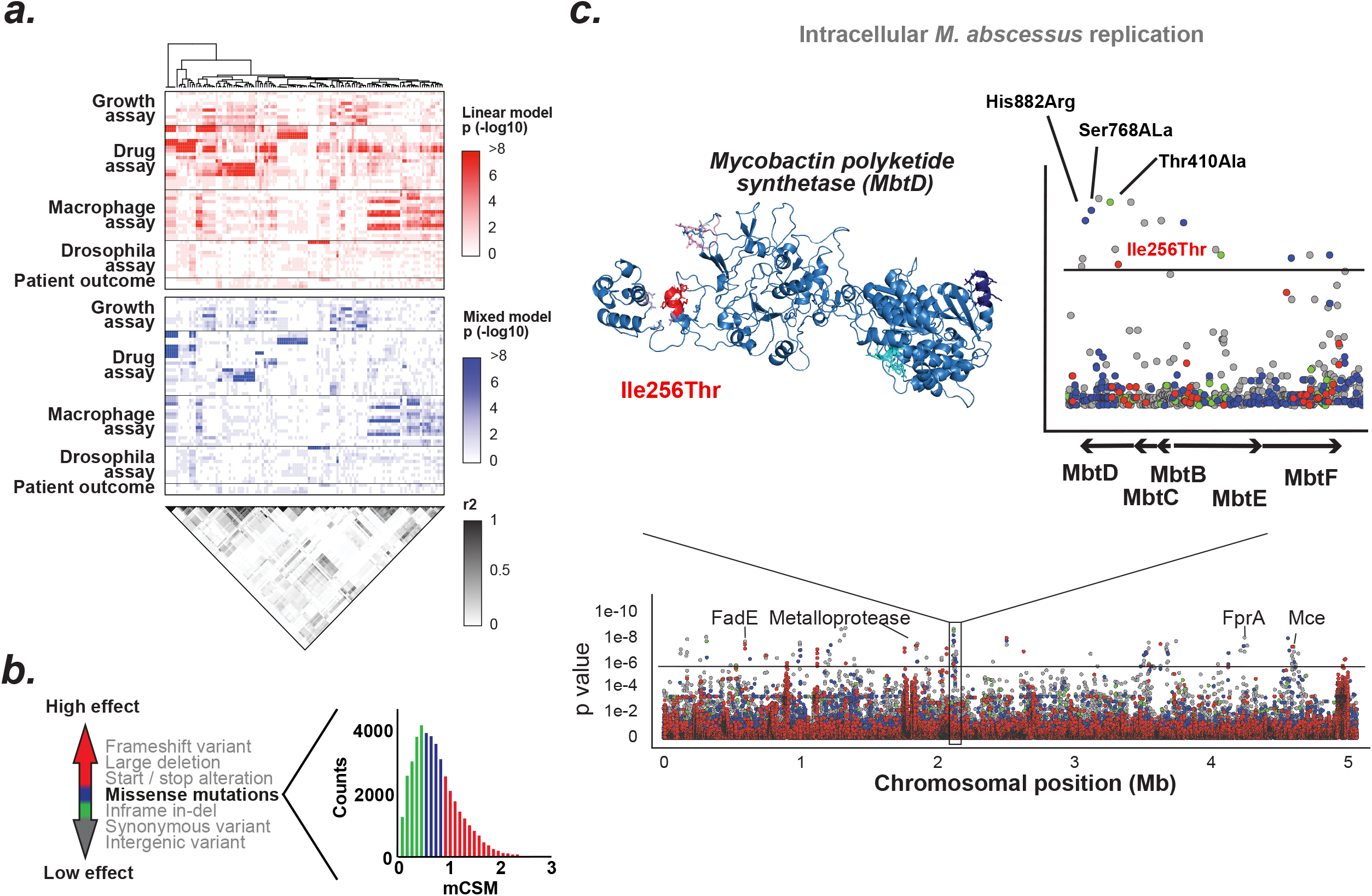
Integrating computational structural modelling into genome-wide association studies. (**a**) Genome-wide associations were performed for all phenotypes with the top variants extracted (up to 5 per association) and ordered using hierarchical clustering (*red* linear model; *blue* mixed model). Pairwise r^2^ measurements of the identified genetic variants (*grey scale*) show extensive genome-wide linkage (LD). (**b**) To identify causal variants and overcome LD, the functional impacts of genetic variants were classified as having high effects (large deletions, frameshifts, start/stop alterations; *red*); moderate effects (inframe insertions/deletions; *blue and green);* and low effects (synonymous and intergenic variants; *grey*). The impact of missense mutations were estimated using proteome-wide computational structural modelling with variants considered as having high (*red*), moderate (*blue*) or low (*green*) functional effects based on terciles of the change in protein stability, estimated using mCSM. (**c**) Manhattan plot of the mixed model GWAS analysis of 264,122 genetic variants for intracellular *M. abscessus* replication. Several loci in *mbtD*, including 4 missense mutations, were identified as potential mechanisms relevant for intracellular *M. abscessus* survival. *Inset*: Protein model of MbtD with the high effect missense mutation Ile256Thr shown in red.

Current GWAS approaches are limited in their ability to accurately identify causal variants by both the presence of linkage disequilibrium, which in the case of *M. abscessus* (as with other bacteria ^28,29^) is extensive and genome-wide (***Figure 2a, Supplementary Figure 6***), and by a failure to consider the impact of mutations on protein function ^30,31^.

We therefore applied proteome-wide computational structural modelling to evaluate the likely functional impact of all nonsynonymous SNPs across the genome, by applying our graphbased machine learning method mCSM ^7^ to our comprehensive *M. abscessus* structural database *Mabellini* ^32^ (***Figure 2b***), in order to identify likely causal mutations.

As an example, the GWAS for intracellular replication of *M. abscessus* within macrophages identified a number of hits at genome-wide significance including a cluster of variants within the mycobactin operon (***Figure 2c***), containing genes involved in iron scavenging ^33,34^. Structural modelling predicted that one variant, a missense mutation (Ile256Thr) in the mycobactin polyketide synthetase (*mbtD*) gene, was most likely to result in loss of protein function and therefore be causally related to the phenotypic change, probably through altering the ability of intracellular *M. abscessus* to access iron.

To understand whether mutations across the genome might have co-evolved, indicating potential epistatic interactions between genes, we deployed correlation-compressed direct coupling analysis (ccDCA ^8^) on whole genome sequences from 2366 clinical isolates of *M. abscessus* to identify whether variant co-occurrence deviated from the expected frequencies based on linkage disequilibrium ^35,36^, and thus indicate evolutionary co-selection. We evaluated 10^12^ potential couplings (resulting from approximately 10^6^ genetic variants), and identified 1,168,913 that were significantly enriched (accepting a false discovery rate of 10^-6^; ***Figure 3a, Supplementary Figure 7***). We found many enriched couplings between known or predicted virulence genes (***Figure 3b, Supplementary Table 2***), indicating pathogenic evolution of *M. abscessus* (as previous identified ^14,37^). We used the ranked outputs from the ccDCA analysis to establish discrete networks of genes that have co-evolved, and thus probably functionally interact (***Figure 3c***). As examples, we find highly connected clusters of: mammalian cell entry (MCE) genes (***Figure 3d***), implicated in controlling adhesion, uptake, and intracellular survival within macrophages ^38,39^; mycobactin synthesis genes (***Figure 3e***), including some identified through GWAS analysis (*Figure 2c,d)*; and genes involved in bacterial secretion systems.

**Figure 3:**
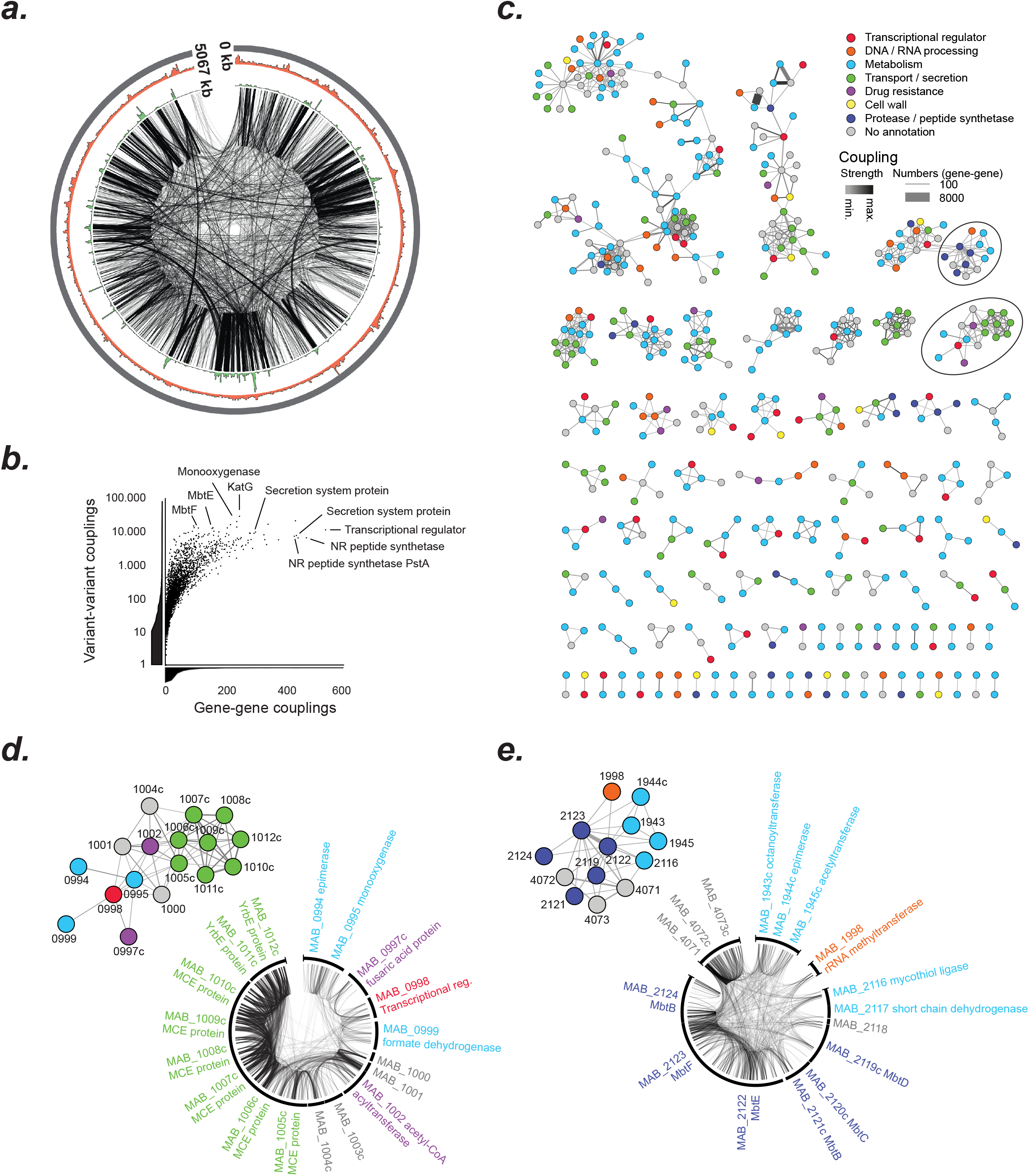
Analysis of genome-wide epistasis through mutational co-evolution. Direct coupling analysis (ccDCA) was used to identify coevolving variants among ~10^12^ potential variant combinations of 2,366 clinical *M. abscessus* isolates. (**a**) Circos plot of the *M. abscessus* chromosome showing the 100,000 top variant-to-variant couplings with a distance of >100bp (*black lines*), coupling density (*green*; range 0-56307 couplings per 5kb) and SNP density (*red*; range 14-1961 SNPs per 5kb). (**b**) Significant variant-variant couplings identified through DCA were pooled as gene-gene couplings and ranked by the number of couplings. (**c**) Networks of co-evolving (and therefore likely epistatic) genes, based on DCA derived genegene couplings, colour coded by functional class. The strength and number of couplings shown by edge colour and thickness respectively. (**d, e**) Examples of highly coupled gene networks, highlighted by circles in (c), involving (d) mammalian cell entry proteins, and (e) components of the mycobactin biosynthesis pathway.

Finally, we sought to integrate outputs from our detailed multidimensional phenotyping, structure-guided GWAS analysis, and DCA-based epistatic mapping, to achieve a systems-level understanding of the genetic basis for important pathological processes in *M. abscessus*.

We focused on *in vivo* infection in *Drosophila*, a model that replicates many of the features of human mycobacterial infection (***Figure 4a***) ^40–43^. Amongst the top hits from our structure-guided GWAS analysis (***Figure 4b, Supplementary Figure 8***) were a deletion in a component of a putative Type II secretion system (*MAB_0471*) and a deleterious mutation in a non-ribosomal peptide synthetase (*MAB_3317c*). Both variants had independently arisen as homoplastic mutations across the *M. abscessus* phylogenetic tree (***Figure 4c***), including within the ancestor of one of the dominant circulating clones (DCC2) of *M. a. abscessus*, responsible for several transmission networks amongst CF patients ^13,14^. We found that isolates with deletions in the Type II secretion system were associated with prolonged survival of infected *Drosophila* and more persistent clinical infection of CF patients (***Figure 4d***).

**Figure 4:**
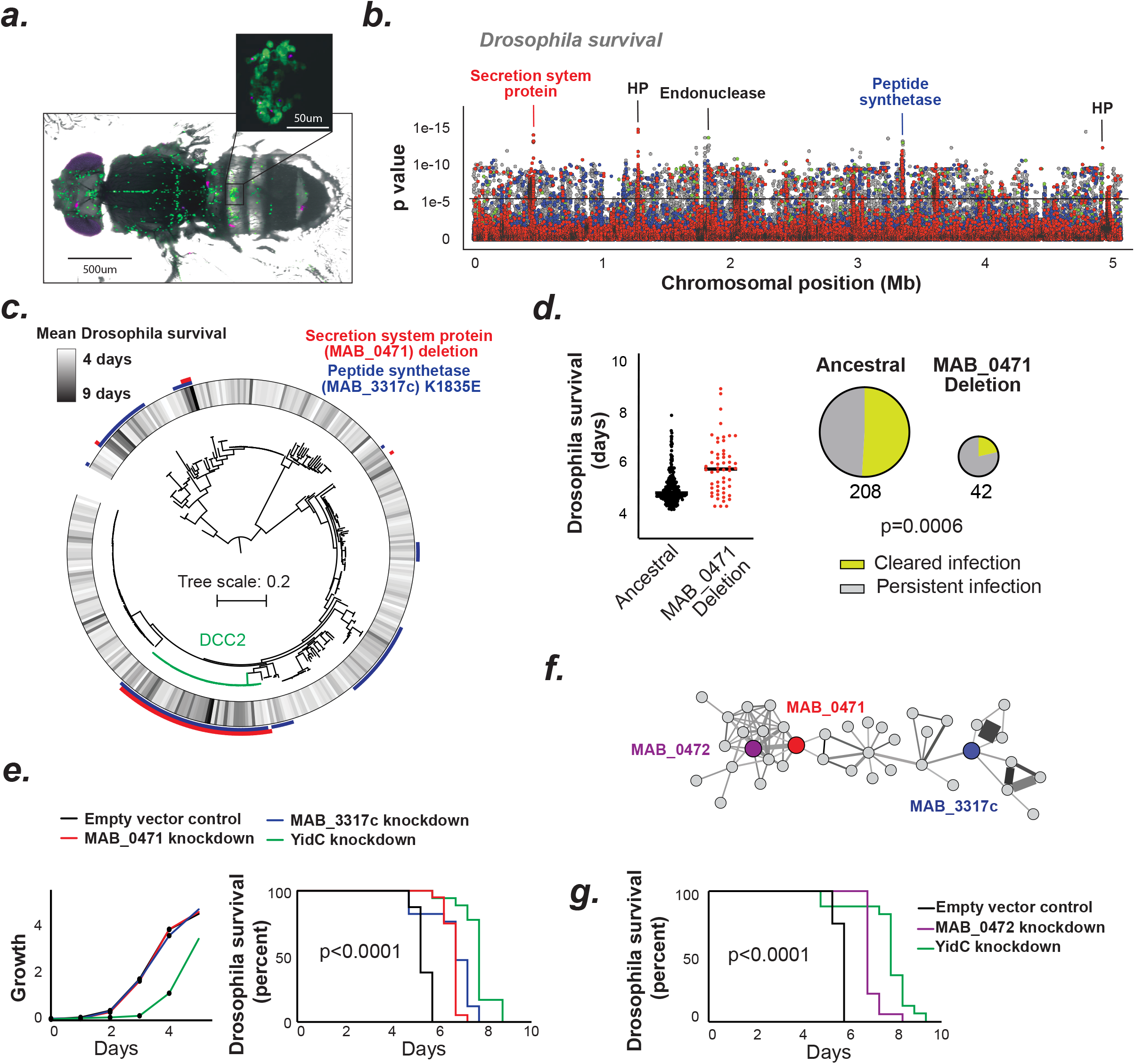
Defining genetic determinants of *in vivo* virulence in *M. abscessus*. (**a**) *Drosophila melanogaster* infected with *M. abscessus (magenta*) resembles mycobacterial infection in other organisms; with infection of phagocytes (*green*) and formation of granuloma-like structures (*inset*). (**b**) Genome-wide association (using a linear model) reveals a putative secretion-system protein and a peptide synthetase to be highly associated with *Drosophila* survival. (**c**) Both variants align to clinical isolates with long survival, including a dominant circulating clone, within the subspecies *M. a. abscessus*. (**d**) Deletion in *MAB_0471* was associated with persistent respiratory infection in cystic fibrosis patients. (e) CRISPR/dCas9 knockdown of *MAB_0471* and *MAB_3317* (unlike the essential gene *yidC*) did not affect growth in liquid culture (*left panel*) but *in vivo* silencing did lead to prolonged survival of infected *Drosophila*, as shown by Kaplan-Meier survival curves generated from data from at least 18 infected flies per bacterial strain. (**f**) Epistatic gene network, derived from DCA outputs, revealed direct coupling of *MAB_0471* with other putative secretion system proteins including *MAB_0472* and a distant connection to the peptide synthetase *MAB_3317*. (**g**) *In vivo* silencing of *MAB_0472* replicated virulence attenuation.

We sought to experimentally validate both these GWAS hits through inducible CRISPR-based transcriptional silencing (CRISPRi) as previously described ^44^. Although we found no effect of gene silencing on growth in liquid media, silencing of either *MAB_0471* or *MAB_3317c* during *in vivo* infection significantly increased *Drosophila* survival (***Figure 4e, Supplementary Figure 9***), indicating that these genes regulate *M. abscessus* virulence.

Our DCA analysis revealed that both these GWAS hits were part of a discrete network of probably epistatic genes involved in bacterial secretion, cell wall biosynthesis, metabolism, and transcriptional regulation (***Figure 4f, Supplementary Figure 10***). To experimentally test this prediction of epistasis, we selected another gene from the same network (*MAB_0472*) and transcriptionally silenced it during *in vivo* infection. We found that *Drosophila* survival was also increased by its CRISPRi knockdown (***Figure 4g***), suggesting that all three genes are functionally interacting.

Thus, phenogenomic analysis can accurately identify critical gene networks responsible for virulence and other characteristics in poorly understood bacterial pathogens such as *M. abscessus*. Our approach of integrating computational structural modelling with conventional GWAS analyses and DCA-driven mapping of gene interaction networks has revealed key determinants of *M. abscessus* antibiotic resistance and virulence.

Importantly we have identified a novel and important gene network, involving a previously unidentified peptide synthetase and type 2 secretion system regulating, that regulates the pathogenic potential of *M. abscessus* and could be targeted therapeutically. Phenogenomic analysis should be readily applicable to any pathogen, permitting rapid identification of prognostic indicators and novel potential drug targets.

## Acknowledgements

We would like to thank John Lees, Philip H. C. Kremer and Simon Harris for statistical and bioinformatical support.

## Author contributions

LB and RAF conceived the project and wrote the manuscript. LB, SB and JS performed the *in vitro* experiments. LB, WHP, KK and MSD performed the *in vivo* experiments. MS and SM performed the computational structural modelling. LB and MS performed direct coupling analysis. LB, AW, IE, JB and BB performed other bioinformatic analyses. SB developed the *M. abscessus* CRISPR interference technique. LB, SB and JS generated bacterial knockdown strains. TLB, MSD, JP and RAF provided supervisory support.

## Ethics approval

Ethical approval was obtained from the National Research Ethics Service (NRES; REC reference: 12/EE/0158) and the National Information Governance Board (NIGB; ECC 3-03 (f)/2012) for centres in England and Wales; from NHS Scotland Multiple Board Caldicott Guardian Approval (NHS Tayside AR/SW) for Scottish centres; and locally for other centres.

## Data availability

All sequencing data of this study is deposited in the European Nucleotide Archive

