## Supplement for "The pathobiology of *Mycobacterium abscessus* revealed through phenogenomic analysis"

#### MATERIALS AND METHODS

##### Sample collection

Samples were obtained from patients with chronic pulmonary disease and respiratory *M. abscessus* infection (as previously reported [1,2]). Isolates were collected in the UK (all major Cystic Fibrosis Centres), the Republic of Ireland (St. Vincent's Hospital Dublin), USA (University of North Carolina Chapel Hill), Sweden (Gotheborg), Denmark (Copenhagen and Skejby), Australia (Queensland) and the Netherlands (Nijmegen). Where possible, *M. abscessus* samples were obtained from the original mycobacterial growth indicator tubes (MGIT), or otherwise from sub-cultured isolates.

##### DNA extraction and whole genome sequencing

*M. abscessus* cultures were sub-cultured on solid media and sweeps of multiple colonies collected for sequencing (as previously described [1,2]). DNA was extracted with the Qiagen QIAamp DNA mini kit. DNA libraries were constructed in pools with unique identifiers for each isolate. Multiplexed paired-end sequencing was performed on the Illumina HiSeq platform.

##### Variant calling

Sequence reads, from 2366 samples, were mapped with BWA to the *Mycobacterium abscessus* reference genome (ATCC19977) followed by an INDEL realignment step using GATK (total alignment) [3–5]. Furthermore, a single random sequence per patient was picked to generate an alignment with a single sample per patient (single patient alignment). Samples with an assembly longer than 6Mb, more than 300 contigs, average depth of coverage below 30x, coverage of the reference genome below 50% or evidence of a mixed infection were discarded. In total, 484 clinical isolates plus the ATCC19977 strain were included in the single patient alignment. Bcftools was used for SNP and small INDEL calling where additional criteria were used to filter SNPs, requiring a minimum base call quality of 50, a minimum mapping quality of 20 and a minimum number of matching reads covering a SNP of 8 (3 per strand) [6]. SNPs were annotated with SNPeff [7]. In addition, regions in the reference genome not or poorly mapped, with a minor allele frequency across all genomes greater than 5% were called as large deletions (gaps): the coverage of 20bp windows with an overlap of 10bp was assessed with sambamba [8]. Two consecutive windows with a mean coverage of 5x or lower (overall mean coverage 75x) were considered a large deletion. If the distribution of consecutive deletions was equal across all isolates these variants were collapsed into a single variant. A maximum likelihood tree of 331 samples assessed for *Drosophila* survival, inferred from SNPs was constructed with RAxML [9].

**Analysis of bacterial growth on different media**

Bacterial growth in nutrient-rich (Middlebrook 7H9 supplemented with 0.4% Glycerol and 10% ADC) or carbon source limited media (Middlebrook 7H9 plus carbon source) was assessed in 96-well plates and quantified with OD600 every 12 or 24h for 10 days. The carbon sources tested were Acetate (10mM), Glucose (2.5mM), Lactate (10mM) and Pyruvate (10mM). Growth of each isolate across all conditions was assessed in quadruplicates. For each well a logistic function was fitted using the R package growthcurver [10]. OD of day 1 was used for early growth and the area under the logistic curve for up to day 10 to assess general growth. The median of the quadruplicates was used as the representative phenotype. If the readout was highly variable (coefficient of variation above 20%) the measurement was considered missing.

**Drug resistance evaluation**

Drug resistance was quantified with minimal inhibitory concentrations (MIC) according to the CLSI guidelines [11]. In brief, ~$5*10^4$ CFUs of each isolate were inoculated in increasing antibiotic concentrations in Mueller Hinton broth (amikacin, cefoxitin, clarithromycin and linezolid) or Middlebrook 7H9 supplemented with 0.4% Glycerol and 10% ADC (clofazimine) per well. Experiments were carried out in duplicates. The MIC was recorded as the lowest drug concentration inhibiting visible growth at days 3, 5, 11 and 14. The mean of both experiments, i.e antibiotic concentration, was recorded and log2 transformed.

**Transformation of clinical isolates**

An expression plasmid carrying tdTomato (obtained from Laurent Kremer) was used to transform clinical isolates, grown in 10ml Middlebrook 7H9 supplemented with 0.4% Glycerol, 10% ADC and 0.05% Tween 80 at 37°C at 100rpm. Competent log-phase bacteria were washed with 10% glycerol containing 0.05% Tween 80. 200ul of the pellet was transferred together with 1ug DNA to a cuvette and electroporated (2500V,1000Ω, 25uF). Transformed bacteria were recovered for 24h in antibiotic-free medium and then transferred to a selective agar plate (7H11 complemented with 10% OADC and 1mg/ml hygromycin). Red colonies were picked and cultured in media containing 1mg/ml hygromycin.

**Generation of single cell suspensions**

The isolates were obtained from frozen stocks and grown in Middlebrook 7H9 (supplemented with 0.4% glycerol, 10% OADC and 0.05 % Tween 80). Exponentially growing isolates were centrifuged at 200g for 5 minutes and the supernatant passed multiple times through a 27-gauge needle before filtrating with a 5um filter (Acrodisc® syringe filter). Single cell suspensions were standardised to a McFarland turbidity of 0.5 and frozen at -80°C.

**Macrophage infection**

THP-1 cells (ATCC TIB-202) were maintained in RPMI 1640 medium supplemented with 10% FCS, Penicillin (100U/ml) and Streptomycin (100U/ml). Around 10.000 THP-1 cells per well were differentiated with 20nM phorbol 12-myristate 13-acetate (PMA) at 37°C in 384-well imaging plates (CellCarrier-384 Ultra, Perkin Elmer). After 2 days the adherent, differentiated THP-1 cells were washed and incubated with DMEM supplemented with 10% FCS. On day 3 post differentiation THP-1 derived macrophages were inoculated with single cell suspensions of clinical *M. abscessus* isolates at a multiplicity of infection (MOI) of 1:5, centrifuged for 10 minutes with 1000 rpm and incubated at 37°C. After 2 hours extracellular cells were washed off. After 2h, 24h or 48h cells were stained with CellMask™ DR (Invitrogen) for 20min, washed, fixed with 4% paraformaldehyde for 1h and stained with DAPI. The cell supernatant was stored at -80°C. The macrophage infection experiments of 245 tdTomato expressing clinical isolates were set up in quadruplicates at once for all timepoints (2h, 24h, 48h).

**High-content image acquisition**

Plates were stored at 4°C and imaged within 24h on the high-content screening platform Opera Phenix® (Perkin Elmer). Spinning disc confocal images of 37 fields per well and 3 fluorescence channels (blue 405/456, red 561/599, far red 640/706) were acquired with a 63x water immersion objective (NA 1.15).

**High-content image analysis**

Automated image analysis was performed with the Columbus™ software (Version 2.9.0, Perkin Elmer). The 37 fields were pooled to single wells. The blue (DAPI) and far-red (CellMask™ DR) fluorescence channels were used to define cells and their borders. A classification algorithm was trained (using supervised machine learning) based on nuclear, cytosolic and cell features to define macrophage viability. Intra- and extracellular mycobacteria were defined using a spot assay on the red fluorescence channel. For each cell as well as the extracellular space the spot area and mean fluorescence intensity was documented. Both measures were used to quantify the mycobacterial load (intracellular load = total sum of [spot area per cell * mean spot intensity per cell]; extracellular load = extracellular spot area * extracellular mean spot intensity; total mycobacterial load = intracellular load + extracellular load). Wells with a cell number of less than 800 were removed; the median of the remaining wells was used. As the most meaningful outputs we reported the fraction of total cells infected (number of *M. abscessus* infected cells / number of total cells), the intracellular and total *M. abscessus* load as well as the fraction of cells alive (number of cells alive / number of total cells). Mycobacterial load or cell kinetics are reflected in the ratio day 2 / day 0 (delta).

**Cytokine assessment**

The supernatant of macrophages was evaluated for IL-8 and TNFα concentrations 24h after mycobacterial infection. TNFα and IL-8 levels were measured in 25µl supernatant on a Luminex 200 instrument (Merck Millipore, UK) using the reagents and protocol supplied with the Milliplex MAP Human Cytokine/Chemokine kit (Merck Millipore, UK).

***Drosophila* infection**

Isogenic flies ($w^{1118}$) were maintained using standard fly medium (2% polenta, 10% Brewer's yeast, 0.8% agar, 8% fructose and water) at 25°C. Flies infected with inducible CRISPRi mutants of *M. abscessus,* were put on tetracycline (0.2mg/ml) supplemented fly medium several days prior to infection. *Drosophila* infections were carried out as reported previously [12–15]. 400 CFUs were injected in 50nl PBS into the abdomen of anaesthetised 6-8 day old male flies. Flies were kept on $CO_2$ for a maximum of 10min, transferred to a new vial and kept at 29°C. Around 15 flies per condition (in total >350 conditions) were infected to assess survival. Fly survival was assessed every 12h until day 10. In order to reduce technical effects related to fly infection, the mean fly survival was calculated excluding the flies dying within the first 3 days and flies where death deviated more than 3 SD from the mean. Fly survival was compared using the log rank test.

**qRT-PCR of *Drosophila* antimicrobial peptides and cytokines**

At least 5 flies were infected with each isolate to assess the immune response to infection. 28 hours after infection, flies were homogenised in 100ul TRIzol (Invitrogen) and stored at -20°C. RNA was then extracted and cDNA synthesis was carried out with the RevertAid Reverse Transcriptase (200U/µl, Thermo Scientific™). qPCRs were performed in duplicates using the Sensimix™ SYBR no-ROX kit (bioline) as reported previously [16,17] with the primers described in in *Supplementary Table 3.*

**Patient outcomes**

Clinical outcome data were available for 300 CF patients (as previously reported [1,2]). Patients were classified as having cleared *M. abscessus* infection (defined as documented culture conversion or a sustained clinical improvement where further cultures were unavailable) or as having persistent infection (if cultures remained positive or the clinical state worsened where no cultures were available) [2]. Lung function decline was estimated as the percentage change in the forced expiratory volume ($FEV_1$) from the available lung function assessment over a period of 12 months from baseline (before infection).

**Phenotype association**

To assess relatedness of phenotypes and phenotypic groups, all phenotype pairs were correlated (Pearson correlation) and a correlation matrix plotted. To identify characteristic phenotypic signatures of clinical isolates, isolates were clustered using representative experimental phenotypes (amikacin MIC d11, clarithromycin MIC d11, growth d10, change in intracellular MAB load, macrophage cell death d2, *Drosophila* attacin level, mean *Drosophila* survival). 199 isolates had at most 1 missing value and were correlated (Person correlation). The resulting correlation matrix was used as a distance measure to cluster isolates with T-distributed stochastic neighbour embedding (tSNE) [18] using the R package Rtsne. Clustering was validated with k-means clustering with a predefined set of 3 clusters (R package kmeans). Phenotypic groups were compared with one-way analysis of variance or chi-squared test, as appropriate; and mapped onto the phylogeny. For each isolate a nearest phylogenetic neighbour was identified; thereby assessing if neighbours are more likely to belong to the identical phenotypic group (chi square of each phenotypic group comparing neighbour pairs vs. non-neighbour pairs)

**Genome-wide association analysis**

Two statistical genome-wide association approaches were employed to assess the effect of individual variants (SNPs, INDELs, large deletions) on phenotypes. A linear mixed model (LMM) controlling for population structure, where the phenotype is modelled on the fixed locus effect and the random effect of the relatedness matrix, was used. However, controlling for population structure considerably reduces power for population-stratified variants [19]. Since population-stratified variants are common in bacteria, genome-wide associations were also analysed with a linear model (LM). Both analyses were performed in GEMMA [20]. The GWAS threshold, i.e. correction for multiple hypothesis testing, was calculated on the effective number of independent high and moderate effect variants. Within the 331 isolates phenotyped for *Drosophila* survival we obtained in total 75260 high/moderate effect variants (large deletions, frameshifts, start/stop alterations, missense mutations) with a minor allele frequency above 0.03. To account for variant dependency due to LD we estimated the effective number of independent tests [21]. 17925 markers were considered independent; after Bonferroni correction we obtained a p-value threshold of $2.8*10^{-6}$. This threshold was applied for all genome-wide associations studies, including those with less variants. Hits were defined as the top 50 significant associations within a phenotype. Manhattan plots were generated using LocusZoom [22].

**Genome-wide protein structure prediction**

As the structures of most proteins in the *M. abscessus* proteome have not been resolved experimentally, it was necessary to model them computationally. We therefore extended our *M. abscessus* structural proteome database, Mabellini [23], which provides only high-confidence, well-annotated structural data, to aim for comprehensive coverage of the entire proteome. Therefore, additional proteins were modelled with lower-confidence templates aided with extensive macromolecular modelling and refinement protocols. The multiple sequence alignments were converted into profile Hidden Markov Models (HMMs), which then have been used to search against a pdb70 (Protein Data Bank chains clustered at 70% sequence identity) database using Hhsearch [24]. The identified templates were then used for comparative modelling, using a modified, MODELLER-based [25], multi-template structure modelling pipeline of Larsson et al. [24]. In addition to structural consensus and an ML-based single-model quality assessment protocol, we also incorporated a rapid method for annotating the quality of protein models through comparison of their distance matrices [26]. As a result, for each of the modelled protein sequences, we obtained a set of theoretical models, ranked by predicted model quality.

**Machine learning for assessing effects of missense mutations**

To evaluate the effect of polymorphisms on *M. abscessus* protein structures, we used the models generated in the previous step to estimate the effect of missense mutations. We applied mCSM (mutation cutoff scanning matrix) [27], which, through graph-based signatures, represents the structural environment of wild type residues and learns which mutations are detrimental to protein structure. For each of the mutations, one or more modelled structures have been used.

**Comparative modelling of *MAB_2119c* (*MbtD*)**

The model of putative polyketide synthase (mbtD, MAB_2119c) was produced as part of Mabellini using the following models: 2hg4, 3tzz, and 2jgp [23]. The Mabellini-derived structure was then subjected to extensive relaxation using Rosetta [28] suite, both in a wild-type and mutated variants, where the lowest energy structure has been chosen for subsequent analysis.

**Ranking of predicted functional impact of SNPs**

Based on SNP annotation (intergenic, synonymous, inframe INDEL, frameshift) and structural modelling predictions of functional impact (see above), variants were allocated to 4 groups: low effect variants (intergenic and synonymous SNPs; *grey*), low-moderate effect variants (inframe INDEL, missense mutations with lowest tertile mCSM scores; *green*), moderate-high effect variants (missense mutations with middle tertile mCSM scores; *blue*) and high effect variants in red (frameshift variant, large deletion, start/stop alteration and missense mutations with highest tertile mCSM scores; *red*).

##### Summary of GWAS hits

To summarise the identified variants across all phenotypes up to 5 significant, highest ranking hits were extracted from each genotype-phenotype association (a single high or moderate effect variant per gene). In total 2x 58 genotype-phenotype associations (LMM and LM) were performed. To assess genetic linkage between these variant hits,  we calculated r2 using PLINK [29].

##### Identification of homologs and construction of multiple sequence alignments

For each of the proteins in the *M. abscessus* proteome, we have constructed a multiple sequence alignment of homologous proteins, which formed a basis for subsequent work. The alignments have been constructed using HHblits, a fast, highly sensitive, HMM-HMM-based sequence search method [30] and used the bundled nr30 database. In the interest of exploring a broader evolutionary landscape of proteins in question, we have decided to include proteins with E-value less than or equal to $10^{-4}$ in the alignment.

##### Genome-wide evolutionary coupling Inference

Exponential models to understand co-evolution in biological sequences have been applied to protein structure prediction [31], and more recently to bacterial genomic sequences. We have previously shown that the method genomeDCA [32] can be effectively employed to understand the co-evolution of *S. pneumoniae* [33], and is extensible and applicable to other systems [33–35]. Here, we employ an approach that blends genomeDCA [32] and CC-DCA [35], to ensure unbiased sampling of evolutionary pressures onto individual positions and pairs of positions across genomic sequences. Correlation-compressed DCA [35] permits genome-wide coupling inference without needing to resort to extensive sampling, as proposed in genomeDCA [32]. We modified this approach to elucidate the effects of low-frequency alleles across the entire *M. abscessus* genome. We conducted at least 60,000 runs, each subsampling 25% of positions in the genome. We defined variant-variant couplings as statistically significant based on the Gumbel distribution (as previously described [21]) corresponding to a false discovery rate of FDR < $10^{-6}$. Variant-variant pairs that spanned a distance of over 100 bp were ranked by coupling strength and visualised plotted on the *M. abscessus* genome using Circos package [36]. Subsequently, we pooled the statistically significant couplings by gene-gene pairs, and ranked them by the number of couplings. Cytoscape was used to plot the network of the 1000 strongest gene-gene couplings, highlighting the number of couplings (edge width), coupling strength (edge colour) and predicted gene function (node colour)) [37].

**Generation of CRISPR interference mutants**

Analogous to CRISPR mediated gene silencing in *Mycobacterium tuberculosis* and *Mycobacterium smegmatis* we established a CRISPR interference platform in *M. abscessus* [38]*. M. abscessus* ATCC 19977 was transformed with pTetInt-dCas9 and a second vector (pGRNAz) containing the small-guide RNA (sgRNA) cassette. For each gene two oligonucleotide were synthesised (forward and reverse), annealed and cloned into pGRNAz. Oligonucletide sequences are outlined in *Supplementary Table 4*. The strains were grown in Middlebrook 7H9 broth (supplemented with 0.4% Glycerol, 10% ADC and 0.05% Tween 80) and selected with hygromycin (1mg/ml) and zeocin (300ul/ml). dCas9 and sgRNA expression were under the control of a tet-inducible promotor. To achieve maximal gene repression cultures were supplemented with 100ng/ml anhydrotetracycline (ATc). As controls an empty vector control and YidC (essential gene) knockdown were used.

### *Supplementary Figure 1*

#### (a) Planktonic growth

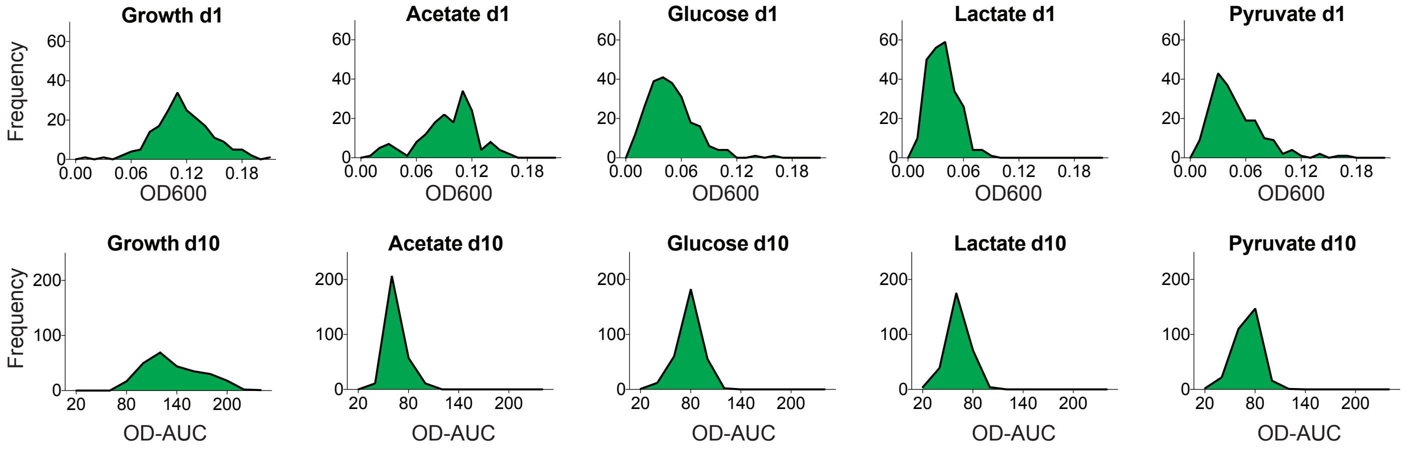


#### (b) Antibiotic resistance

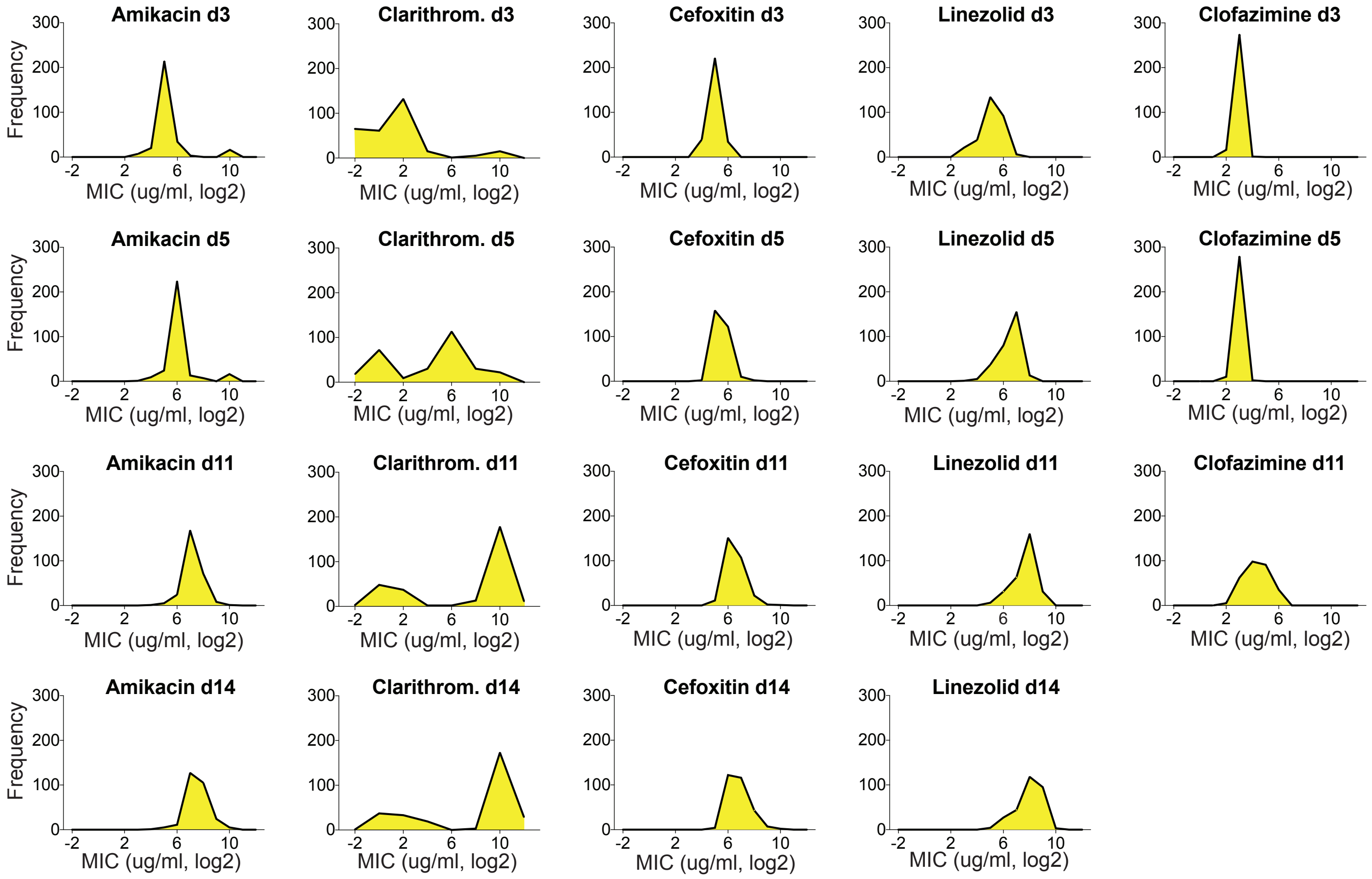


#### (c) Clinical outcome

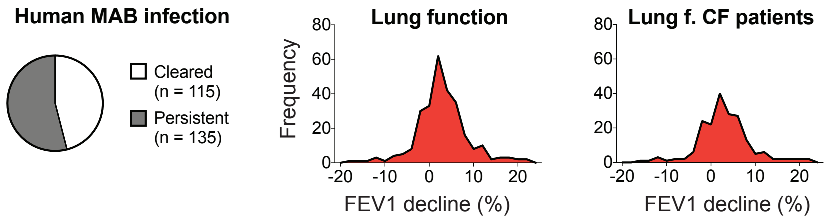

#### (d) Macrophage infection

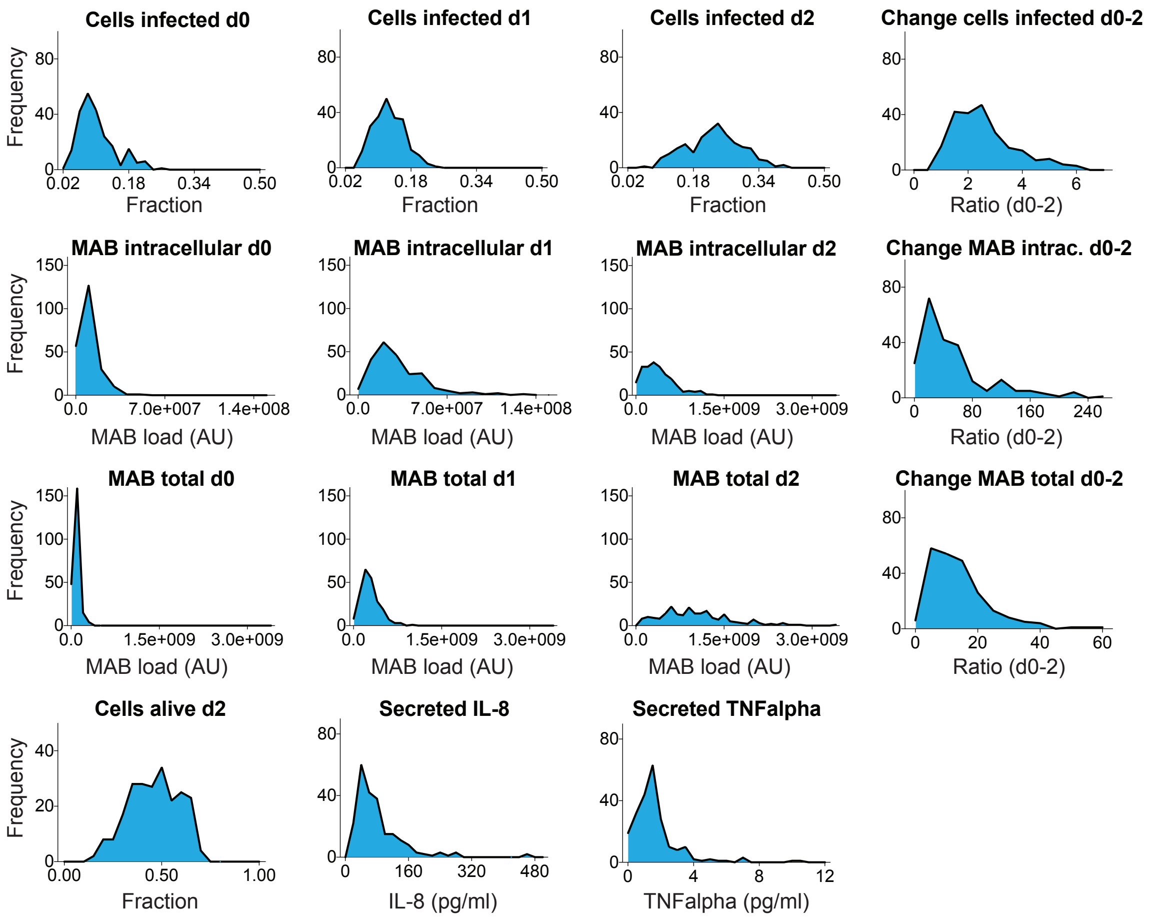


#### (e) *In vivo* infection

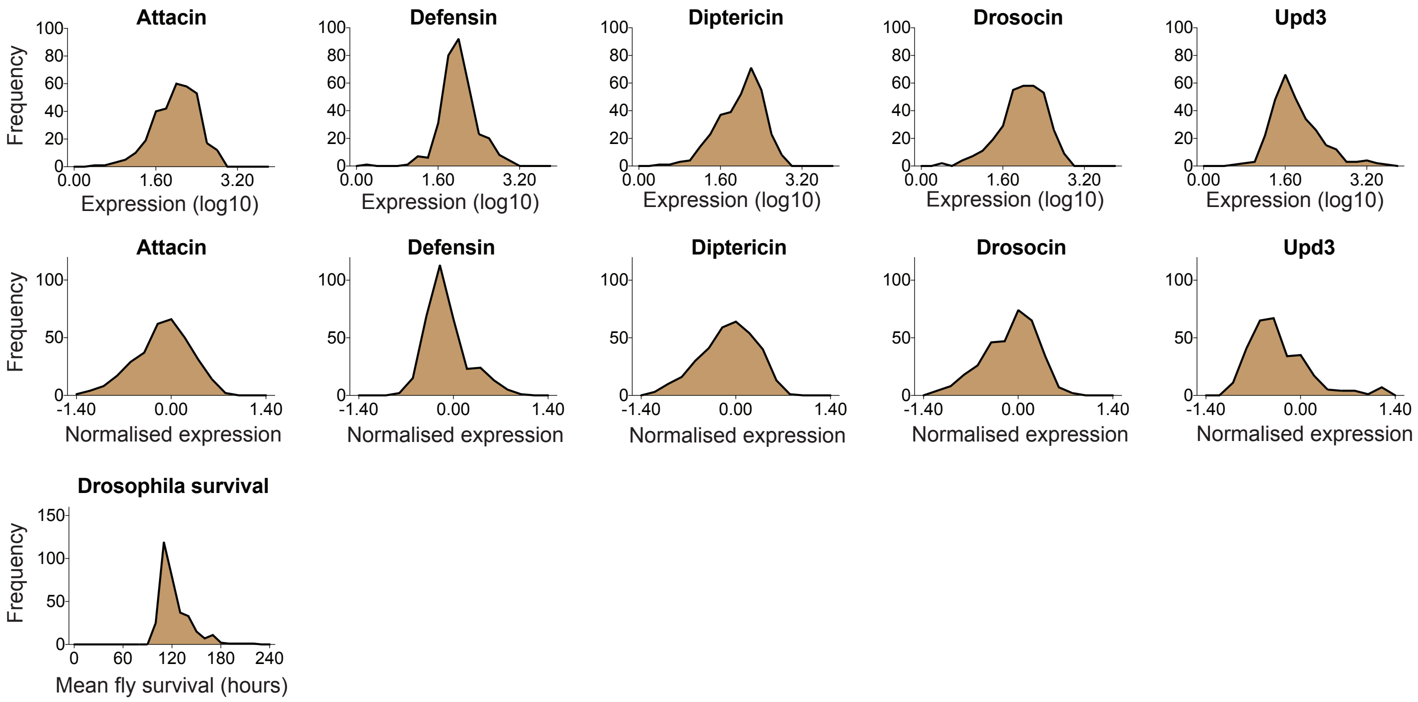


**Figure S1. Distribution of phenotypic behaviour of *M. abscessus* clinical isolates** for the following phenotypes: **(a)** Planktonic growth in different carbon sources; (b) Antibiotic susceptibility; (c) Clinical outcomes; (d) Macrophage infection; (e) *In vivo* infection

### *Supplementary Figure 2*

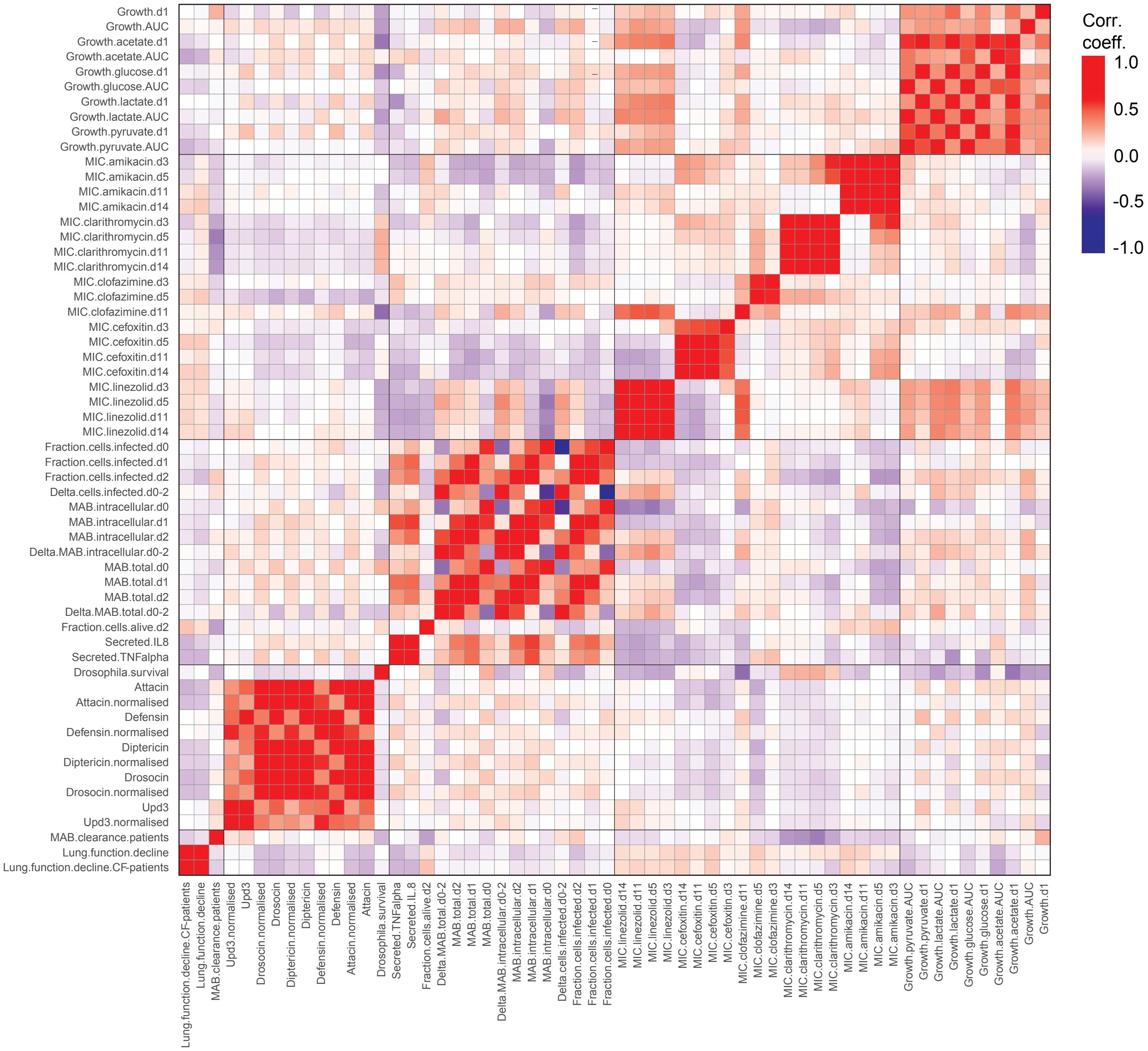


**Figure S2. Pearson correlation matrix of and *in-vitro* and *in-vivo* bacterial phenotypes and clinical outcome.**

#### *Supplementary Figure 3*

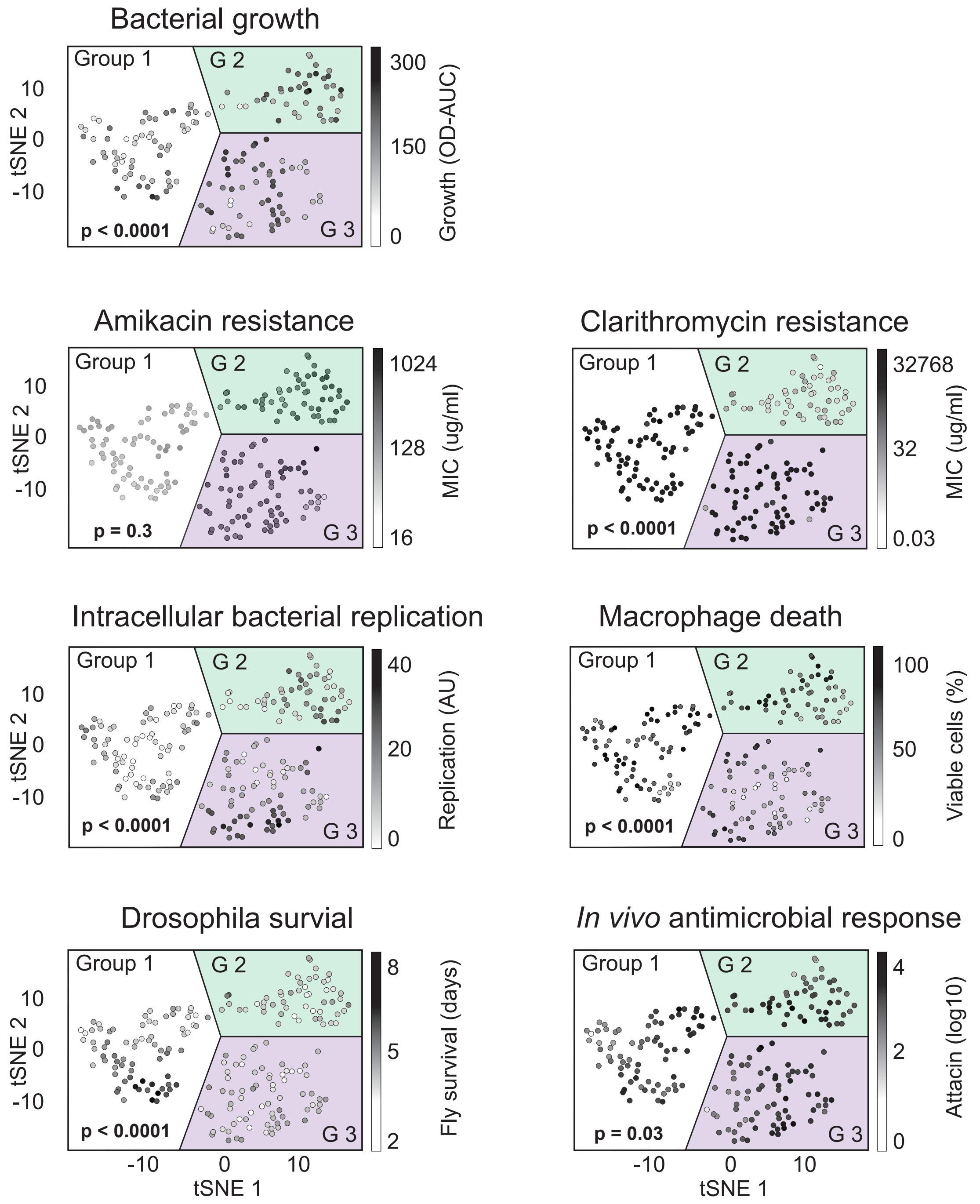


**Figure S3. Characteristics of phenotypic clusters.**
The 7 phenotypes bacterial growth, drug resistance (amikacin and clarithromycin), intracellular replication, macrophage death, Drosophila survival and antimicrobial response were used to group clinical *M. abscessus* isolates. tSNE plots highlight the cluster allocation of respective phenotypes. The three clusters were compared using the one-way analysis of variance.

### *Supplementary Figure 4*

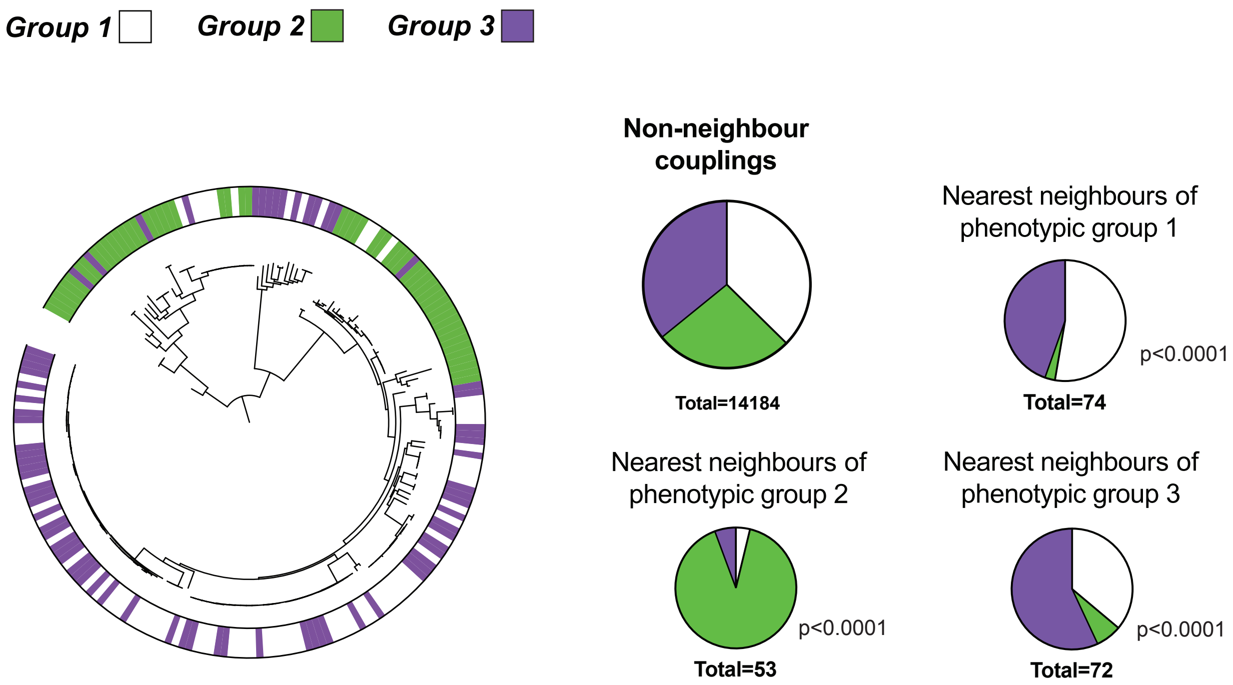


**Figure S4. Mapping of phenotypic groups to phylogeny.** Maximum likelyhood tree of 199 isolates and their phenotypic groups. Pie charts of nearest phylogenetic neighbours of respective groups. Distributions of groups were compared against a random distribution (non-neighbour couplings) using the chi-squared test.

### *Supplementary Figure 5*

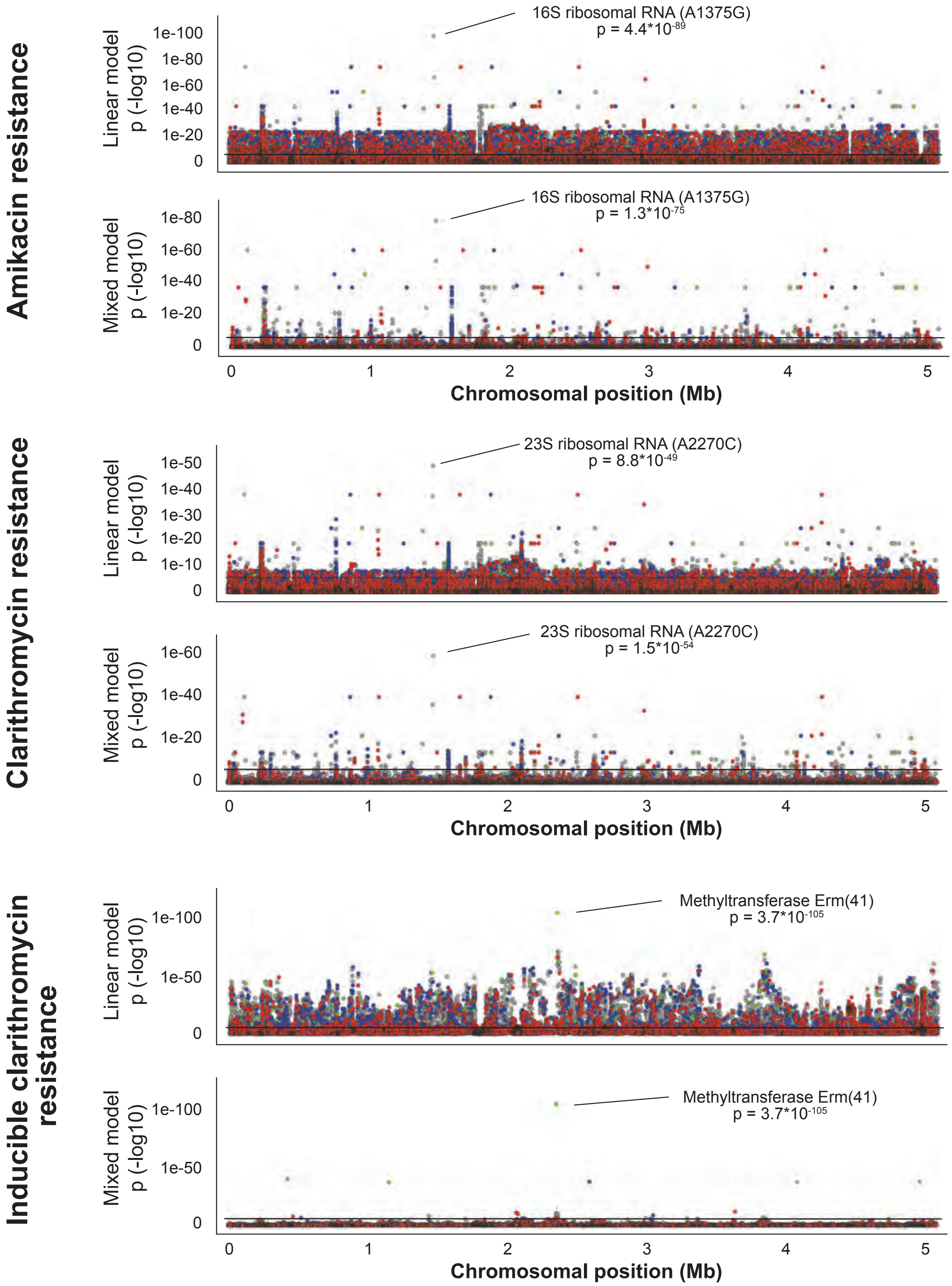


**Figure S5. Genome-wide association of known resistance mechanisms.** Genotype-phenotype associations of amikacin and clarithromycin MICs Day 3 revealed the known resistance loci in the 16S and 23S ribosomal RNA, respectively. Similarly, *erm(41)* conferring inducible macrolide resistance, was identfied when assessing clarithromycin MICs at Day 11 in *M. abscessus subsp. abscessus*.

### *Supplementary Figure 6*

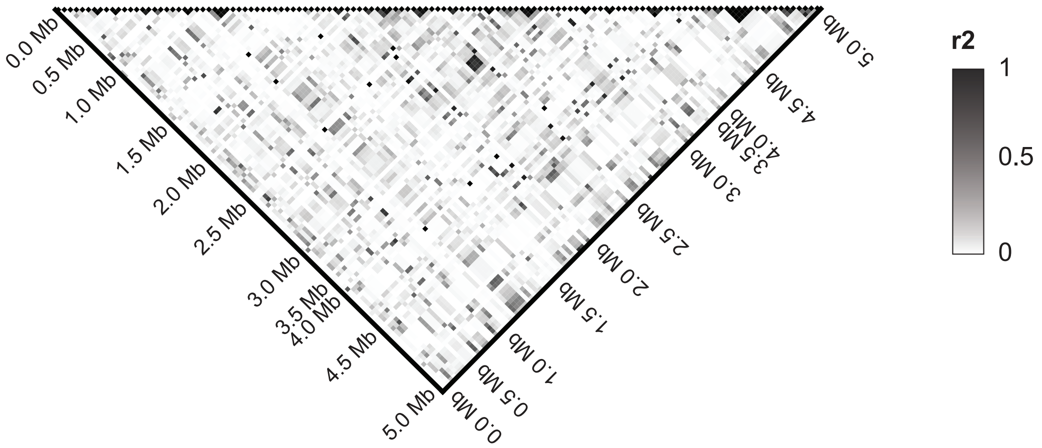


**Figure S6. Pairwise r2 measurements of variants ordered by genomic position.**

### *Supplementary Figure 7*

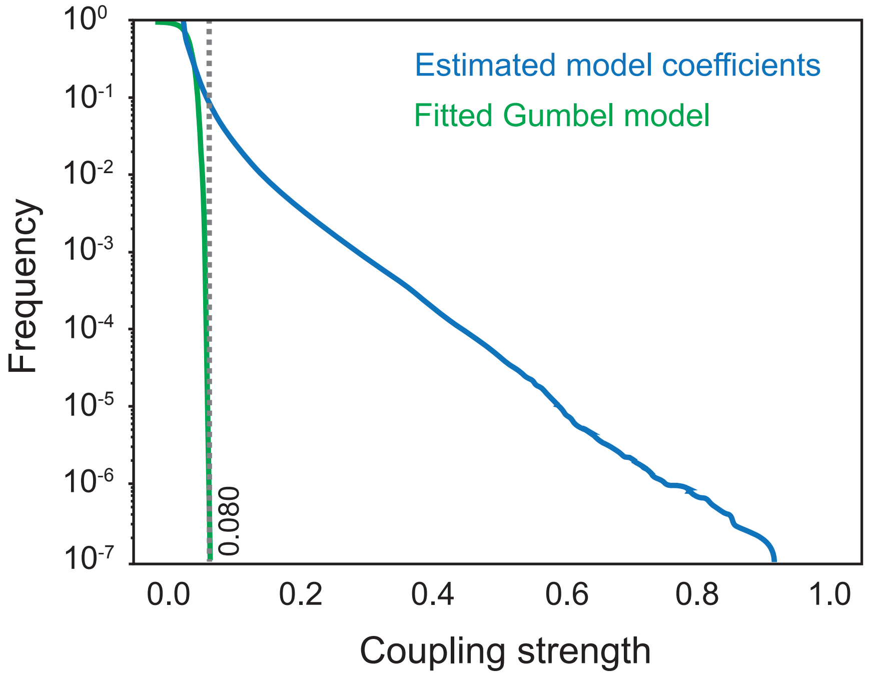


**Figure S7. Distribution and threshold of the variant-variant coupling strength in *M. abscessus.*** Divergence of theoretical (fitted distribution) and empirical distribution of the coupling strength. The dashed line highlights the defined threshold (coupling strength above 0.080) with a false discovery rate of 1 in $10^6$ couplings.

#### *Supplementary Figure 8*

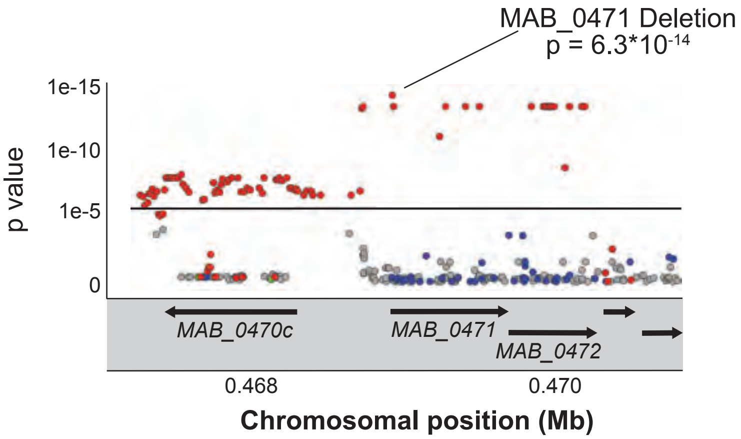


**Figure S8. Genetic variants around MAB_0471 associated with *Drosophila* survival.**

### *Supplementary Figure 9*

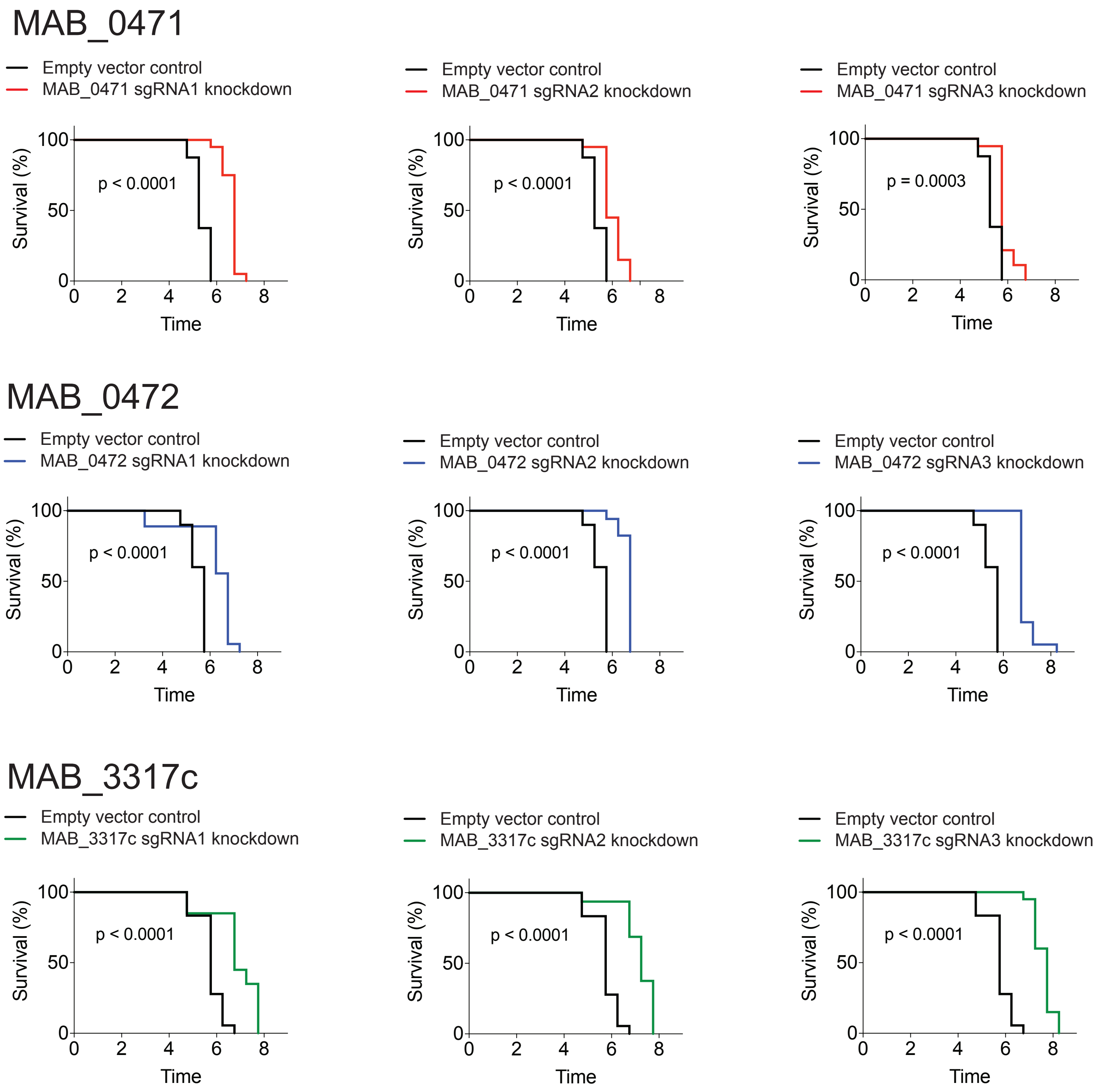


**Figure S9. CRISPR/dCas9 knockdown of target genes using different guide RNAs**

#### *Supplementary Figure 10*

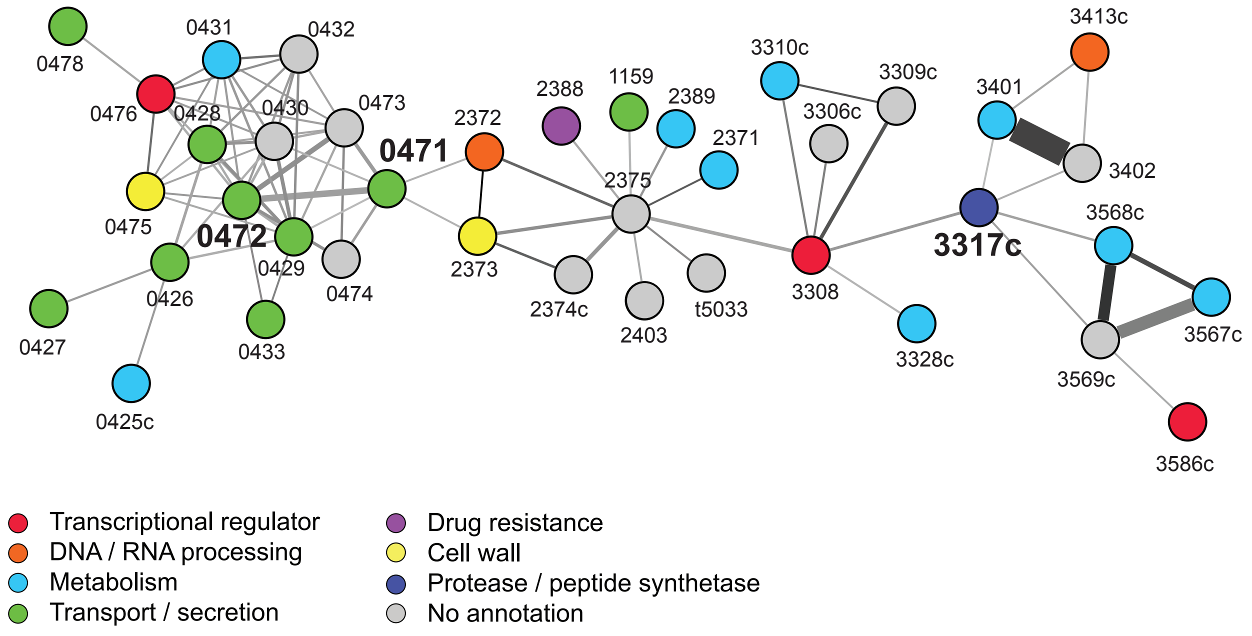


**Figure S10. Epistatic interactions of MAB_0471 and MAB_3317.**
